# Increasing the Throughput and Reproducibility of Activity-Based Proteome Profiling Studies with Hyperplexing and Intelligent Data Acquisition

**DOI:** 10.1101/2023.09.13.557589

**Authors:** Hanna G. Budayeva, Taylur P. Ma, Shuai Wang, Meena Choi, Christopher M. Rose

## Abstract

Intelligent data acquisition (IDA) strategies, such as real-time database search (RTS), have improved the depth of proteome coverage for experiments that utilize isobaric labels and gas phase purification techniques (i.e., SPS-MS3). While most applications of IDA have been focused on the analysis of protein abundance, these approaches have recently been applied to activity-based proteome profiling (ABPP) studies aimed at characterization of protein site engagement by small molecules. In this work, we extend IDA capabilities offered by vendor software through a program called InSeqAPI. First, we demonstrate robust performance of InSeqAPI in the analysis of biotinylated cysteine peptides from ABPP experiments. Then, we describe PairQuant, a method within InSeqAPI designed for the hyperplexing approach that utilizes protein-level isotopic labeling and peptide-level TMT labeling. PairQuant allows for TMT analysis of 36 conditions in a single sample and achieves ∼98% coverage of both peptide pair partners in a hyperplexed experiment as well as a 40% improvement in the number of quantified cysteine sites compared to non-RTS acquisition. We applied this method in ABPP study of ligandable cysteine sites in the nucleus leading to an identification of additional druggable sites on protein-and DNA-interaction domains of transcription regulators and on nuclear ubiquitin ligases.

## Introduction

Isobaric mass tags are widely utilized in proteomics research for relative quantification of protein levels and post-translational modifications. For example, tandem mass tag (TMT) reagents allow for analysis of up to 18 conditions in a single mass spectrometry (MS) run.^1^ Gas phase purification through synchronous precursor selection MS3 (SPS-MS3) data acquisition improves the accuracy of isobaric label based quantitation, but can decrease proteome depth due to a longer duty cycle as compared to Orbitrap MS2 analysis alone.^2^ To improve the depth of SPS-MS3 experiments, a real-time database search (RTS) has been introduced to enable online identification of peptides from ion trap MS2 spectra and limit collection of SPS-MS3 scans to instances where a peptide is confidently identified.^3,4^ This approach increases the amount of instrument duty cycle dedicated to collecting quantitative data associated with identified peptides and results in an increased number of proteins quantified by SPS-MS3.^3,4^

Recently, RTS-aided acquisition of TMT data was applied in activity-based proteome profiling (ABPP) studies for characterization of accessible and reactive cysteine residues.^5^ Cysteine is the most reactive nucleophilic residue in the proteome and cysteine residues are present in the active sites of many enzyme families (such as kinases BTK^6^ and EGFR^7^) and near ligandable pockets of “undruggable” proteins (such as KRAS G12C^8^). Their identification and characterization may be used in the discovery of new covalent drug targets (reviewed in^9^). There are approximately 214,000 cysteine sites encoded in the human genome. Combined results from a few recent cysteine profiling studies revealed ∼10,000 ligandable cysteine sites against ∼300 electrophilic compounds of various reactivities.^5,10,11^

While the number of available TMT reagents continues to increase, currently analyzing more than 18 samples (including various cell types, treatments, and replicates) requires performing multiple TMT experiments and combining data through one or more common bridge channels.^12^ Due to the stochastic sampling of data dependent acquisition, comparing quantitative data across multiple TMT experiments leads to an increase of missing data as peptides and proteins are not quantified in all multiplexed experiments.^13^ This is particularly true for site-level post translational modification (PTM) data which requires peptides containing the PTM to be identified across multiple TMT experiments.^14^

It was previously demonstrated that TMT tags can be combined with stable isotope labeling in cell culture (SILAC) in a “hyperplexing” approach to assess temporal abundance of yeast proteins or to profile site-resolved protein turnover.^15,16^ Hyperplexing increases the number of samples that can be quantified in a single experiment, but also increases sample complexity potentially decreasing the total number of unique peptides that are quantified. Hyperplexed data acquisition also utilizes data-dependent acquisition and it is not guaranteed that both peptides within a hyperplexed pair will be chosen for identification and quantification. To date, these challenges have limited the broad adoption of hyperplexing in quantitative proteomics experiments.

In this study, we present an approach that combines protein-level isotopic labeling (by SILAC or by isotopic cysteine-reactive probes) and peptide-level quantification by TMT to perform hyperplexed ABPP experiments. First, we introduce inSeqAPI - an instrument application interface (iAPI) program that enables creation of custom intelligent data acquisition (IDA) methods, including those that utilize a real-time database search. To address the challenges of hyperplexed experiments we describe a novel IDA method called PairQuant that utilizes a real-time peptide identification to improve the quantitation of both peptides within hyperplexed pairs. Lastly, we utilize PairQuant to perform a 36-plex cysteine reactivity profiling experiment in isolated nuclei treated with a set of electrophilic fragment compounds to investigate ligandability of nuclear targets.

## Materials and Methods

### Cell lines

C2C12 mouse myoblasts were obtained from ATCC and maintained in Dulbecco’s Modified Eagle Medium (DMEM) supplemented with 10% FBS and Glutamax. MCF7 adenocarcinoma cell line was purchased from ATCC and maintained in RPMI-1640, 10% FBS and Glutamax.

For SILAC experiments, C2C12 cells lines were grown in DMEM SILAC “light” ([^12^C,^14^N] Lys and [^12^C,^14^N] Arg) or SILAC “heavy” ([^13^C,^15^N] Lys and [^13^C,^15^N] Arg) media for five passages, until >95% incorporation was achieved.

### Reagents

Light and heavy desthiobiotin iodoacetamide (LDBIA and HDBIA) reagents were synthesized by Pharmaron and stored at 4.92mg/mL and 5mg/mL in DMSO, respectively. Structures of the DBIA reagents are provided in **Supp. Figure 1**. IodoTMTsixplex^TM^ reagent (Thermo Fisher Scientific) was reconstituted in methanol prior to the experiment, according to the manufacturer’s protocol. Electrophilic compound fragments were synthesized in-house (G35, G59, G84, G92), purchased from Sigma (KB05), or Enamine (G51, KB02, KB03) and stored in DMSO at 50mM concentration.

### Compound reactivity profiling (cysteine trapping assay)

Incubation was carried out in a 100 mM phosphate buffer (pH 7.4) at 37°C with gentle shaking. Test compounds (1 µM), diphenhydramine (0.1 µM, pre-internal standard), and cysteine (5 mM) were mixed sequentially to start the reaction. Aliquots were taken from 0, 30, 60, 90, 120, 150, and 180 minutes and quenched with 1x volume of ice-cold acetonitrile containing propranolol (2 µM, post-internal standard). The resulting samples were quantified via LC-MS/MS. Percentage remaining was calculated using peak area ratios normalized to the 0 minutes time-point sample. Half-life and k_react_ were then determined from fitting the data with one phase decay model (% *remaining* = 100 ∗ *e*^−*kreact* ∗ 5 *mM* ∗ *time*^). Incubation without cysteine was also carried out to check compound stability in the buffer.

### Sample processing for activity-based proteome profiling workflow

For benchmarking an RTS against a non-RTS method in ABPP analysis with DBIA probes, C2C12 mouse myoblasts were collected from cell culture by scraping, washed with PBS and frozen until later or immediately lysed. Lysis was performed on ice in PBS/1% NP-40 buffer and lysates were pulse-sonicated with a probe. Insoluble fraction was removed by 4°C centrifugation at 12000xg for 10 minutes and protein concentration in the soluble fraction was measured by Pierce^TM^ BCA protein assay kit (Thermo Fisher Scientific). Protein concentration was adjusted to 1-2 mg/mL for LDBIA treatment.

For whole-cell proteome profiling with electrophilic fragments, lysates were treated with indicated compounds at 500µM concentration for 1 hr at room temperature. Unreacted cysteine sites were labeled with 500µM LDBIA reactive probe for 1hr at room temperature. Upon labeling, protein mixtures were reduced with 5mM Pierce^TM^ DTT (Thermo Fisher Scientific), 30 minutes at room temperature and alkylated with 20mM Pierce^TM^ iodoacetamide (Thermo Fisher Scientific) for 30 minutes in the dark at room temperature. Protein mixtures were cleaned up by chloroform-methanol precipitation. In brief, four volumes of methanol, one volume of chloroform and three volumes of water were added to the mixture and centrifuged for 2 minutes at 20,000xg. Top layer was removed and the remaining protein disk was washed twice with methanol. After final wash, all liquid was removed and the protein pellet was air dried for 15 minutes.

Protein pellets were redissolved in 20mM HEPES, pH 8.0 and digested for 4 h at 37°C with lysC (1:50 enzyme:protein ratio), followed by overnight digestion at 37°C with trypsin (1:100 ratio). TMTPro^TM^ (Thermo Fisher Scientific) labeling was performed as described in the manufacturer’s protocol. TMT tag incorporation of >95% and relative protein abundances between channels were checked by MS prior to quenching with 5% hydroxylamine and mixing at 1:1 ratio.

Streptavidin enrichment was performed on AssayMap Bravo (Agilent) automation using affinity enrichment protocol. Briefly, lyophilized TMT-labeled peptide mixtures were resuspended in PBS. Streptavidin cartridges (5uL capacity) were equilibrated with PBS and loaded with the samples. Upon loading, cartridges were washed sequentially with PBS, 0.1% PBS and HPLC water. Biotinylated peptides were eluted with 50% acetonitrile/0.1% TFA. Eluents were dried down prior and desalted by C18 PhyTip columns (Biotage) using PhyNexus MEA robot (Biotage) prior to LC-MS analysis.

For iodoTMT experiment, MCF cell pellets were lysed with HENS buffer (Thermo Scientific) by pipette mixing before sonicating in a cold water bath, followed by 10 minute incubation on ice. Protein lysate obtained by high speed centrifugation, 14,000xg at 4°C for 15 minutes, and separation of the protein solute from the pelleted cellular debris was collected. Protein concentration was brought to 1 mg/mL and reduced with a final concentration of 10 mM TCEP for 1 hour at 50°C. IodoTMT reagent was prepared as per manufacturer’s protocol (Thermo Scientific) and labeling was carried out for 1 hour at 37°C in the dark. Labeling was quenched with a final concentration of 20 mM DTT (Sigma), incubating at 37°C in the dark for 15 minutes. Samples labeled with iodoTMT were combined after quenching. Protein cleanup was performed via a methanol chloroform precipitation as described above. Subsequently, the pellet was dissolved with 2 M UREA (in 20 mM HEPES), sonicated before adding trypsin (Promega) at a 1:60 ratio of enzyme/substrate and digested overnight at 37°C. Following digestion, peptides were acidified with 1% TFA before desalting with a 3 cc C18 Sep-Pak columns (Waters, Milford, MA) eluting with 60% ACN/0.1% TFA. Samples were lyophilized in preparation for enrichment.

For affinity enrichment of iodoTMT-labeled peptides, anti-TMT resin slurry (Prod#90076, ThermoFisher Scientific) was washed 3 times with TBS. Peptides were then re-suspended with TBS before addition to the rinsed resin. Enrichment was performed by incubating for 2 hours at room temperature with end-over-end mixing. The supernatant was removed and the resin washed 4 times with 2 M UREA in TBS, followed by 4 times with just TBS, and 3 times with water. Peptides were eluted off the beads with 50% ACN/0.5% TFA after incubation for 10 minutes with shaking and elution was performed twice. All centrifugation steps were performed at 2,500 xg for 1 minute at room temperature. After drying to completion, peptides were cleaned up using a 5 µl C18 Phytip (Biotage, Inc) and injected for LC-MS/MS acquisition.

### Mass Spectrometry analysis for ABPP workflow

For DBIA labeling experiment, desalted samples were reconstituted in 2% acetonitrile/0.1% formic acid (buffer A) at ∼1ug/uL peptide concentration for liquid chromatography-tandem mass spectrometry (LC-MS) analysis on Dionex Ultimate 3000 RSLCnano system (Thermo Fisher Scientific, Inc) and Orbitrap Eclipse Tribrid MS (Thermo Fisher Scientific, Inc). Peptides (∼1ug) were resolved on 25 cm x 75 µm Aurora column packed with 1.6 µm C18 (Ion Opticks) by 135 minutes linear gradient of 4% to 30% buffer B (98% acetonitrile/0.1% formic acid) in buffer A (2% acetonitrile/0.1% formic acid) at a flow rate of 300 nL/min.

Where indicated, standard SPS-MS3/Multi-Notch-MS3 TMT method^2^ was performed with the following parameters. Each duty cycle included an MS1 scan in the Orbitrap at 120,000 resolution across 350-1350 m/z range with normalized automatic gain control (AGC) target of 250%, 50 ms maximum injection time. Data dependent ion trap MS2 scans were performed on the top 10 peptides with CID activation, 0.5 m/z isolation window in quadrupole, turbo scan rate, 150% normalized AGC target, 35 ms maximum injection time. Eight MS2 fragment ions were selected for Orbitrap SPS-MS3 scans with isolation widths of 2 m/z using isolation waveforms with multiple frequency notches. These ions were fragmented during MS3 by high energy collision-induced dissociation (HCD) and analyzed by Orbitrap at 50,000 resolution, 400% normalized AGC target, 150 ms maximum injection time.

For LDBIA-based experiments benchmarking inSeqAPI method against Thermo RTS and standard SPS-MS3 method, MS1 and MS2 scans were acquired with the same settings as standard SPS-MS3 TMT method. For Thermo RTS and inSeqAPI RTS analysis of mouse cell lines, the following parameters were used: Uniprot mouse database October 2022 version, including 25440 entries of common contaminants, Swissprot sequences of canonical and protein isoforms, plus decoys. Static modifications included Cys carbamidomethylation (+57.0215), K and n-term TMTZero^TM^Pro (+295.1896); variable modifications included Met oxidation (+15.9949), Cys with light desthiobiotin label (+239.1633). Other parameters were set to 0 maximum missed cleavages, 3 variable modification per peptide, FDR filtering enabled, 35 ms maximum search time. Comet scoring threshold: 0.5 Xcorr, 0.05 dCn, 10 ppm precursor mass error, +2 charge state. InSeqAPI modification node allowed for inclusion of only biotinylated peptide for MS3 analysis. MS3 HCD settings were as in standard SPS-MS3 mode, except 600% normalized AGC target and 400 ms maximum injection time.

For iodoTMT sample analysis, data was collected on an Orbitrap Eclipse Tribrid (Thermo Fisher Scientific, San Jose, CA USA). Peptides loaded onto a IonOptick Aurora GEN3 C18 column (1.7 µM, 75 µM x 25 cm) via a Nanospray Flex Ion-Source (Thermo Scientific) at a voltage of 1.6 kV with a flow rate of 0.4 µL/min in Solvent A (98% water/2% acetonitrile/0.1% formic acid). Separation was achieved at a flow rate of 0.3 µL/min with a linear gradient of 5% solvent B (98% acetonitrile/2% water/0.1% formic acid) to 35% solvent B over 158 minutes with a total run time of 180 minutes. For SPSMS^3^ acquisition, full ms scans were collected in the orbitrap at 120,000 resolution, scan ranging from 350 to 1600 m/z, automatic gain targeting (AGC) of 1 x 10^6^, and a maximum injection time of 50 ms. MS^2^ ions were selected using a top speed data dependent mode and fragmented with CID energy of 35, AGC of 1.5 x 10^4^, and a maximum injection time of 100 ms. Utilizing the Orbitrap Fusion Tribrid’s SPS mode for isolating MS^3^ ions, the top 8 MS^2^ precursor ions are selected and fragmented via higher collision dissociation energy (HCD) of 55, AGC of 7.5 x 10^4^, a maximum injection time of 350 ms, isolation width of 1.2 Da, and a resolution of 50,000 at 200 m/z. For ThermoRTS acquisition all parameters matched that of the SPSMS3 method with the exception of an increased AGC 1 x 10^5^ target for MS3 ions and the addition of Thermo’s real-time search module enabled. Real time search parameters include a human proteome with a decoy addendum from the October 2022 version database for tryptic peptides (42374 total entries). Variable modifications of methionine oxidation (+15.9949 Da), and variable modifications of cysteine (+329.2266) for IodoTMT. Other parameters include, allowance of 1 missed cleavage, 1 maximum variable mod/peptide, enabling FDR filtering, TMT SPS MS3 Mode and a maximum search time of 35 ms. Scoring Thresholds include 0.5 Xcorr, 0.05 dCN, 10 ppm of the precursor and a 2+ charge state.

InSeqAPI method acquisition functions by enabling Xcalibur method triggering of MS1 and MS2 levels of which parameters match that of the SPSMS3 method indicated above, whilst all MS3 specific parameters are controlled via the API.

### Sample processing for activity-based proteome profiling workflow with hyperplexing

For optimization of RTS acquisition of hyperplexed samples, two alternative workflows were tested. In HLDBIA/TMT hyperplexing workflow, MCF7 lysates at 2 mg/mL protein concentration were labeled with 500 µM LDBIA or HDBIA reagent, as described in regular ABPP workflow above. For hyperplexed samples with SILAC labeling, “heavy” SILAC or “light” SILAC protein mixtures from C2C12 lysate at 2 mg/mL concentrations were labeled with 500 µM LDBIA. For both types of hyperplexed samples, “heavy” and “light” protein mixtures were reduced, alkylated, and combined 1:1 into a single sample (HLDBIA or SILAC). Peptide mixtures were cleaned by chloroform/methanol precipitation and digested with lysC/trypsin mixture at 1:50 enzyme:protein ratio as described above. Peptides were labeled with TMTPro^TM^Zero following the manufacturer’s protocol. Prior to quenching, TMT labeling efficiency was checked to ensure >95% labeling. Streptavidin enrichment was performed on AssayMap Bravo, as described above.

To fractionate hyperplexed samples, eppendorf tips were loaded with SDB-XC Empore disks and activated with 100% acetonitrile, followed by 2 washes with 20 mM NH_4_HCO_2_. Lyophilized peptides were resuspended in 20 mM NH_4_HCO_2_ and loaded onto the disks, followed by two washes, first elution with 12% acetonitrile/20 mM NH_4_HCO_2_ and second elution with 50% acetonitrile/20 mM NH_4_HCO_2_. Elutions were dried down prior to injection.

### Nuclear lysate treatment

For nuclear fractionation, MCF7 cells were propagated to six 15 cm plates and collected by scraping into ice-cold PBS when at ∼90% confluency. Nuclear extraction was performed using REAP protocol. ^17^ Briefly, cell pellets were resuspended in 0.1% NP40 and triturated 5 times. After 10 sec pulse-spin, the supernatant (cytosol) was removed, followed by an additional wash with 0.1% NP40. Pelleted nuclei were resuspended in PBS and sonicated on tip with thirty 1 sec pulses, 20% AMP. Insoluble fraction was removed by centrifugation at 14,000xg for 10 minutes at 4°C. Soluble nuclear fraction was treated with indicated compounds at 100 µM for 1hr at room temperature. Unlabeled reactive cysteine sites were probed with 500 µM LDBIA or HDBIA for 1 hr at room temperature. Samples were reduced with 5 mM DTT for 30 minutes at room temperature and LDBIA/HDBIA samples pairs were combined 1:1. Combined samples were alkylated with 20 mM iodoacetamide for 30 minutes at room temperature. Sample clean-up, digestion and TMTPro labeling was performed as described above.

### Mass Spectrometry analysis for ABPP workflow with hyperplexing

Hyperplexed samples were reconstituted in buffer A for liquid chromatography-tandem mass spectrometry (LC-MS) analysis on Dionex Ultimate 3000 RSLCnano system (Thermo Fisher Scientific, Inc) and Orbitrap Eclipse Tribrid MS (Thermo Fisher Scientific, Inc). Peptides (∼1 ug) were resolved on 25 cm x 75 µm Aurora column packed with 1.6 µm C18 (Ion Opticks) by 135 minutes linear gradient of 4% to 30% buffer B (98% acetonitrile/0.1% formic acid) in buffer A (2% acetonitrile/0.1% formic acid) at a flow rate of 300 nL/min.

Where indicated, standard SPS-MS3/Multi-Notch-MS3 TMT method^2^ was performed with the same parameters as in regular ABPP workflow described above. In addition, a “Targeted Mass Difference” node was used for acquisition of precursor isotopic peptide pairs with the following parameters: for SILAC experiment, delta M1 was set to 8.0142 and 10.0082 and for HLDBIA experiment, delta M1 was set to 6.0319; partner intensity range set to 10-100%; subsequent scans performed on both ions in the pair, with the same charge state.

For SILAC/TMT hyperplexed sample analysis using Thermo RTS and InSeqAPI RTS acquisition, the following RTS parameters were used: Uniprot mouse database October 2022 version, including 25,367 Swissprot sequences of canonical and protein isoforms, plus common contaminants and decoys; static modifications included Cys carbamidomethylation (+57.0215), K and n-term TMTZero^TM^Pro (+295.1896); variable modifications included Met oxidation (+15.9949), Cys with light desthiobiotin label (+239.1633), Lys +8.0142, Arg +10.00827. Other parameters were set to 0 missed cleavages, 3 variable modifications per peptide, FDR filtering disabled, 35 ms maximum search time, and protein filter set to exclude decoy matches from MS3 analysis. Comet scoring threshold: 0.5 Xcorr, 0.05 dCn, 10 ppm precursor mass error, +2 charge state. MS3 HCD settings were as in standard SPS-MS3 mode, with 600% normalized AGC target and 400 ms maximum injection time.

For “heavy” SILAC/TMT sample analysis, the following RTS parameters were used: Uniprot mouse database (October 2022 version, including 25,367 Swissprot sequences of canonical and protein isoforms, plus common contaminants and decoys; static modifications included Cys carbamidomethylation (+57.0215), n-term TMTZero^TM^Pro (+295.1896), Lys TMTZero^TM^Pro +295.1896, Arg +10.00827; variable modifications included Met oxidation (+15.9949), Cys with light desthiobiotin label (+239.1633). Other parameters were set to 0 missed cleavages, 3 variable modifications per peptide, FDR filtering enabled, 35 ms maximum search time. Comet scoring threshold: 0.5 Xcorr, 0.05 dCn, 10 ppm precursor mass error, +2 charge state. MS3 HCD settings were as in standard SPS-MS3 mode, with 600% normalized AGC target and 400 ms maximum injection time.

For “light” SILAC/TMT sample analysis, the following RTS parameters were used: Uniprot mouse database October 2022 version, including 25,367 Swissprot sequences of canonical and protein isoforms, plus common contaminants and decoys; static modifications included Cys carbamidomethylation (+57.0215), Lys and n-term TMTZero^TM^Pro (+295.1896); variable modifications included Met oxidation (+15.9949), Cys with light desthiobiotin label (+239.1633). Other parameters were matched to “heavy” SILAC/TMTPro sample analysis.

For HLDBIA/TMT hyperplexed sample analysis using Thermo RTS and InSeqAPI RTS acquisition, the following parameters were used for RTS: Uniprot human database (October 2022 version, includes 42,352 Swissprot sequences of canonical and protein isoforms, plus decoys); static modifications: Cys carbamidomethylation (+57.0215), K and n-term TMTZero^TM^Pro (+295.1896); variable modifications: Met oxidation (+15.9949), Cys with light desthiobiotin label (+239.1633) and Cys with heavy desthiobiotin label (+245.1952). Other parameters were set to 1 maximum missed cleavages, 2 max variable modification per peptide, FDR filtering disabled, 35 ms maximum search time, and protein filter set to exclude decoy entries from MS3 analysis. Comet scoring threshold: 0.5 Xcorr, 0.05 dCn, 10 ppm precursor mass error, +2 charge state. MS3 HCD settings were as in standard SPS-MS3 mode, except 600% normalized AGC target and 400 ms maximum injection time.

For LDBIA/TMT and HDBIA/TMT sample analysis using Thermo RTS and InSeqAPI RTS acquisition, the following parameters were used for RTS: Uniprot human database October 2022 version (includes 42,352 Swissprot sequences of canonical and protein isoforms, plus decoys); static modifications included Cys carbamidomethylation (+57.0215), K and n-term TMTProZero (+295.1896); variable modifications included Met oxidation (+15.9949) and Cys with light desthiobiotin label (+239.1633) or Cys with heavy desthiobiotin label (+245.1952). FDR filtering was enabled, whereas other parameters were matched to the hyperplexed sample analysis.

### InSeqAPI Instrument Control

inSeqAPI was written in C# (.Net Framework 4.6.1) and interfaces with Orbitrap Tune versions 3.4 and higher on Orbitrap Eclipse and Ascend mass spectrometers. Search functionality within inSeqAPI is enabled by a previously described version of Comet (v.2020013, comet-ms.sourceforge.net/).^4^ Real-time access to spectral data was enabled by the Thermo Scientific instrument API (https://github.com/thermofisherlsms/iapi). For inSeqAPI methods, the native instrument method was used to control the nano-LC as well as MS1 scans, filters guiding selection of precursor ions for MS2, and MS2 scans (as described above). inSeqAPI received the MS2 scans collected by the instrument and analysis was triggered by specific MS2 scan descriptions.

The inSeqAPI ‘real-time search’ method contained a single node comprising a real-time search filter to match the spectra to a peptide sequence, PPM adjustment filter to account for systematic mass shifts, an LDA filter to perform real-time false discovery estimations, a decoy filter to ensure decoy identifications are not quantified, a modification filter to only quantify peptides that contained a modified cysteine residue, a mass range filter to remove y1 ions, a precursor range exclusion filter to exclude peaks around the precursor, a tag loss exclusion filter to exclude fragment ions that lost the TMT tag (i.e., TMTc ions), a targeted SPS ion filter to only select fragment ions that contain the quantitative tag, and finally a topN filter to select the most abundant fragment ions for SPS-MS3 analysis. If all of these filters were passed, instructions for an SPS-MS3 scan were sent to the instrument.

The inSeqAPI ‘PairQuant’ method contained two nodes: 1) PairQuant node, 2) PairSeq node (**Supp. Fig. 6**). The PairQuant node comprised the same filters as the ‘real-time search’ method described above with the addition of a PairQuant filter that keeps a record of which peptides have been identified and saves spectral information relating to which SPS ions would be selected for SPS-MS3 and a PairSeq filter that keeps a record of which peptides have been identified. If all filters are passed the PairQuant node will send instructions for two SPS-MS3 scans corresponding to the light and heavy partners of the hyperplexed pair. The PairSeq node contains only a PairSeq filter (the same PairSeqFilter object that is referenced in the PairQuant node) that checks to see if both peptides within a hyperplexed pair have been identified. The ‘Targeted Mass Difference’ filter within the vendor method isolates and fragments each peptide within a pair in successive scans. Due to this the PairSeq filter assumes that both peptides within a hyperplexed pair will be sequenced within 2 scans. If only one peptide is sequenced, the PairSeq node will send instructions to perform an ion trap MS2 on the missing peptide in an attempt to sequence it. This MS2 will be analyzed within the PairQuant node just as MS2 scans which are triggered by the vendor method.

### Data analysis for ABPP workflow

Offline search was performed using comet v.2019.01 with parameters matched to the RTS search. Linear Discriminator Algorithm was used for peptide filtering to <2% FDR, as previously described.^18^ The PSMs were filtered out if they were from peptides with length less than 7. For quantification of site ligandability, total reporter ion intensity across all 18 channels / plex was filtered to >10,000 and individual reporter ion intensity less than 2^8 noise estimate. In the case of redundant PSMs (i.e. multiple PSMs in one MS run corresponding to the same peptide ion), only the single PSM with the least missing values or highest isolation specificity or highest maximal reporter ion intensity was retained for subsequent analysis. If the same PSM was identified in multiple fractions, only the single PSM with the highest isolation specificity or least missing values or highest sum of reporter ion intensity were kept. The reporter ion intensities of all the peptide ions mapped to a cysteine site were summarized into a single site level intensity in each channel and plex by MSstats (v4.8.3).^19^ For each cysteine site, the local normalization on the summaries was performed to reduce the systematic bias between plexes. For the local normalization, we created an artifact reference channel by averaging over all the 18 channels for each cysteine site and plex. The cysteine site abundances in the reference channel of each plex were equalized to their median across two plexes. MSstatsTMT R package (v2.8.0)^20^ estimated log2(fold change) and the standard error by linear mixed effect model for each cysteine site. The inference procedure was adjusted by applying an empirical Bayes shrinkage. To test two-sided null hypothesis of no changes in abundance, the model-based test statistics were compared to the Student t-test distribution with the degrees of freedom appropriate for each site and each dataset. The resulting P values were adjusted to control the FDR with the method by Benjamini–Hochberg.

For iodoTMT analysis, MS/MS spectra were searched using comet against Uniprot human database October 2022 version (includes 42,352 Swissprot sequences of canonical and protein isoforms, plus decoys). Search parameters included trypsin cleavage with allowance of up to 2 missed cleavage events, a precursor ion tolerance of 50 ppm, and a fragment ion tolerance of 0.8 Da. Searches permitted variable modifications of methionine oxidation (+15.9949 Da), and variable modifications of cysteine (+329.2266) for iodoTMT. Reporter ions produced by the IodoTMT tags were quantified with an in-house software package known as Mojave. MSstatsTMT R package was used for statistical analysis as described above.

### Gene ontology enrichment analysis

A list of Uniprot accession entries identified in the nuclear ABPP experiment were uploaded to DAVID.^21,22^ UniprotKB Keywords Cellular Component enrichment analysis was performed against proteins identified in the whole-cell ABPP experiment as a background.

### Data and Software availability

Mass spectrometry raw data files, metadata, quantification data and statistical analysis results have been uploaded to the UCSD MassIVE mass spectrometry data repository (https://massive.ucsd.edu/ProteoSAFe/static/massive.jsp) using identifier MSV000092113 and password “hyperplexing”. The inSeqAPI software enabling inSeqAPI RTS and PairQuant data collection is available upon request and requires users to have signed the Thermo IAPI agreement as well as a distribution agreement with Genentech, Inc.

## Results

### Development of intelligent data acquisition approaches for TMT profiling of reactive cysteine sites

In a typical activity-based proteome profiling study of ligandable cysteine sites, cells or native cell lysates are treated with an electrophilic compound that reacts with ligandable sites (**Figure 1A**). Subsequent incubation with a reactive probe, such as desthiobiotin-modified iodoacetamide (DBIA), labels all of the remaining reactive sites. For relative quantification between conditions, peptide mixtures are labeled with tandem mass tags (TMT), mixed and biotinylated peptides are enriched by streptavidin. Site ligandability can be quantified based on the relative abundance of peptides containing DBIA labeled cysteine residues in compound-treated and vehicle-treated samples.

**Fig 1.**
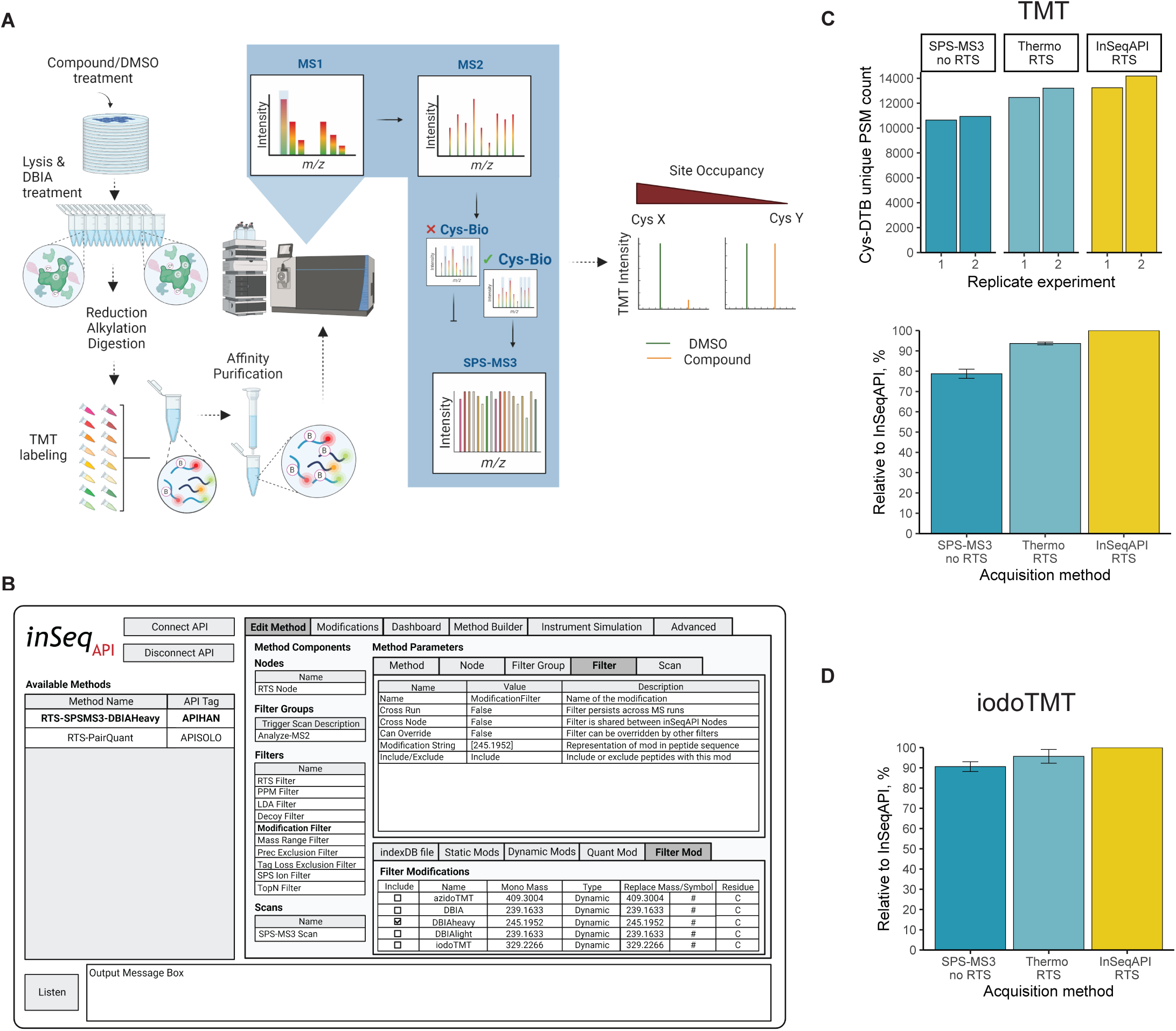
Development and evaluation of InSeqAPI intelligent data acquisition platform in activity-based proteome profiling workflow. **A**. Schematic of activity-based proteome profiling workflow with real-time database searching and MS3 acquisition of biotinylated cysteine peptides. **B**. User interface of InSeqAPI, an intelligent data acquisition application. **C**. C2C12 mouse myoblast lysates were treated with 500 µM LDBIA probe for 1 hr. Equal amounts of enriched cys-containing biotinylated peptides were injected for analysis by SPS-MS3 without RTS, with vendor RTS and InSeqAPI RTS methods in two biological replicates. Number of unique cys-DTB peptide-spectrum matches and their relative values compared to InSeqAPI RTS are compared (top and bottom barplot, respectively, n=2). **D**. MCF7 cell lysates were labeled with iodoTMT reagent for 1hr and TMT-labeled peptides were enriched with anti-TMT resin. Enriched peptide mixture was analyzed with RTS or without RTS, as indicated (n=2).

Real-time search (RTS) MS data acquisition methods that greatly improve the depth of TMT-based quantification studies were recently introduced within the vendor software for select instruments (Orbitrap Eclipse and Orbitrap Ascend).^3,4^ In this approach, ion trap MS2 spectra containing precursor fragment ions, are collected by the instrument and searched in real time against a database of tryptic peptides. Only spectra matched to a peptide with high confidence are sent for acquisition of Orbitrap SPS-MS3 spectra that record relative abundances of TMT report ions (**Figure 1A**). This approach improves the depth of SPS-MS3 analysis by limiting the collection of slow SPS-MS3 scans to peptides that have been identified and dedicating a larger percentage of instrument duty cycle to the collection of SPS-MS3 scans that will be used in the final quantitative analysis. However, these intelligent data acquisition approaches provided by instrument vendors have been optimized for measurements of protein abundance. These methods typically make assumptions that may not be applicable to experiments quantifying PTM bearing peptides. For example quantitative scans are triggered for all peptide identifications, even if users only want to quantify peptides that contain a particular PTM. This is an important consideration for reactive cysteine profiling where we wished to limit collection of quantitative scans to peptides that contained DBIA labeled cysteine residues.

Due to these limitations, we set out to tailor the RTS acquisition method to the analysis of ABPP samples through the development of an intelligent data acquisition (IDA) instrument application program interface (iAPI) called inSeqAPI (**Figure 1B**). Like other iAPI programs^3,4^, inSeqAPI captures data that are collected by the mass spectrometer in real-time, analyzes the data through a number of user-defined filters, and creates instructions for new scans that are sent back to the mass spectrometer for execution. The differentiating characteristic of inSeqAPI is the ability to develop customized methods through modular method design that reflects the construction of methods within the vendor software. Methods within inSeqAPI are broken down into separate nodes that each contain a list of filters that are successively applied to the spectra (**Figure 1B**). If all filters are passed the instructions to perform the scan(s) specified within the node are sent to the instrument. Importantly, a method can contain multiple nodes, enabling a unique set of filters to be applied to different types of scans as indicated by the scan descriptions. Methods can be easily edited by re-ordering filters and changing filter parameters within the graphical user interface (GUI) (**Figure 1B**). Lastly, new filters are implemented through the creation of a single C# class, quickly enabling new functionality such as the ‘Modification Filter’ used to ensure only identified peptides that contain a modified cysteine residue are quantified (**Figure 1B**).

To benchmark InSeqAPI against non-RTS and vendor RTS (Thermo RTS) acquisition methods for ABPP studies, native cell lysates of C2C12 myoblasts were treated with “light” DBIA (LDBIA) reactive probe, followed by TMT labeling and streptavidin enrichment of biotinylated peptides. For MS data acquisition, all MS1, MS2 and MS3 parameters were matched for accurate comparison of the methods. However, it was observed that the RTS acquisition method sampled more low-intensity precursor ions than the non-RTS method (**Supp. Figure 2A**). Therefore, MS3 AGC target and Max IT were optimized in RTS acquisition approach to boost TMT intensity and signal/noise (**Supp. Figure 2B**). Overall, a 15% increase in the number of quantified biotinylated cysteine peptides was observed with Thermo RTS acquisition compared to non-RTS method (**Figure 1C**). Furthermore, an additional 7% increase was yielded by the InSeqAPI RTS method. A boost in the number of quantified cysteine-containing peptides by InSeqAPI RTS was largely attributed to a greater number of total spectra acquired (**Supp. Figure 2C**). We further tested InSeqAPI performance in iodoTMT cysteine labeling approach. As expected, we observed that RTS approaches allow for deeper proteome coverage compared to a non-RTS method (**Fig. 1D**). Overall, our data demonstrates that InSeqAPI generates comparable or better results than vendor software and therefore can be utilized in real-time database search of mass spectrometry data.

### Improving hyperplex pair quantification with PairQuant within inSeqAPI

Due to the stochastic nature of data-dependent MS acquisition methods, quantifying peptides or proteins across multiple TMT plexes can lead to missing data. For instance, we observed a loss of ∼40% of replicate peptide quantification data when comparing the number of Cys-DTB (desthiobiotin-labeled cysteine) peptide pairs quantified between two TMT runs (**Supp Fig. 3**). In order to improve the number of replicate quantification between TMT plexes, we employed two alternative workflows utilizing protein-level isotopic labeling. In the first workflow, SILAC is used for labeling of cells in replicate samples, followed by lysis, labeling with DBIA probe, mixing of SILAC replicates and processing them as a single plex for TMT-based quantification (**Fig. 2A**). In the second workflow, replicate samples are labeled with isotopically “heavy” DBIA (HDBIA) and “light” DBIA (LDBIA), mixed and also processed as a single plex for TMT-based quantification (**Fig. 2A, DBIA structures in Supp. Fig. 1**). Therefore, in each workflow, a hyperplexed sample is formed containing protein-level isotopic tags for replicate analysis and peptide-level TMT tags for comparisons between conditions.

**Fig 2.**
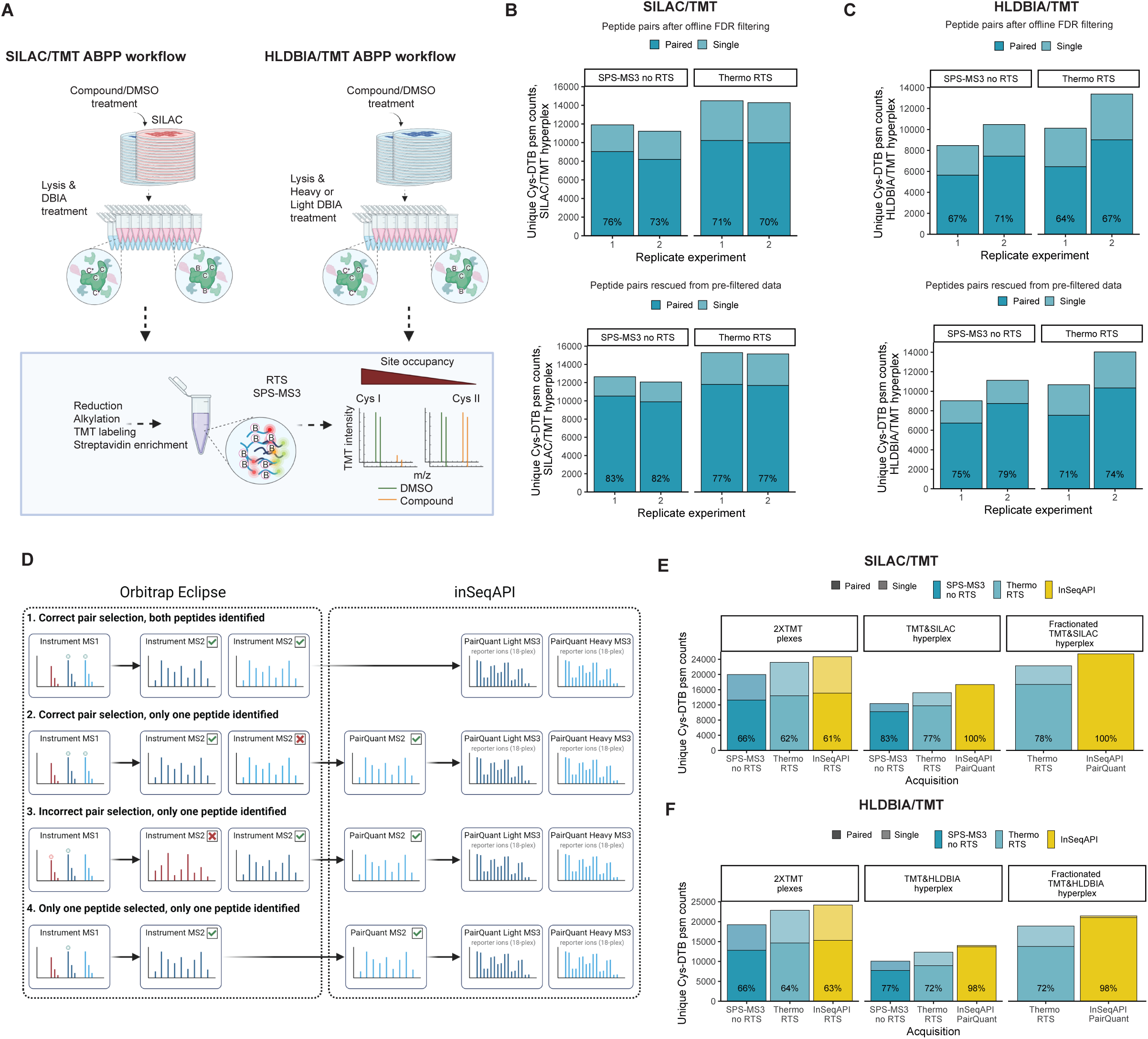
Sample hyperplexing in activity-based proteome profiling. **A.** Workflow schematic for activity-based proteome profiling with sample hyperplexing. In SILAC/TMT hyperplex, isotopic labeling is done on protein level by “heavy” or “light” SILAC, followed by isobaric TMT labeling on peptide level. In HLDBIA/TMT hyperplex, isotopic labeling on protein level is done by “heavy” or “light” DBIA, followed by TMT labeling on peptide level. Duing MS analysis, Targeted Mass Difference node is enabled for detection of isotopic peptide pairs in MS1. **B and C.** C2C12 SILAC lysates were labeled with 500 µM LDBIA probe for 1 hr (B) or MCF7 lysates were labeled with 500 µM HDBIA (“heavy”) or LDBIA (“light”) probe for 1 hr (C). Number of isotopically labeled biotinylated PSM pairs (both replicates acquired) and PSMs without pairs (missing replicate measurement) is plotted for each experiment. Top panel shows PSMs present after offline FDR filtering. Bottom panel includes additional PSMs rescued from unfiltered data due to the presence of their isotopic pairs in the filtered data. Data from two individual experiments is shown. **D**. PairQuant acquisition of hyperplexed peptide pairs within InSeqAPI for SPS-MS3 analysis. (1) Both hyperplexed peptides within pair are fragmented and identified triggering SPS-MS3. (2) Both hyperplexed peptides within pair are fragmented but only one is identified, triggering PairQuant MS2 to identify missing peptide and trigger SPS-MS3. (3) Incorrect selection of hyperplexed peptides leading to only one peptide identified prompts PairQuant MS2 to identify missing peptide and trigger SPS-MS3. (4) Only one peptide within pair is fragmented prompting PairQuant MS2 to identify missing peptide and trigger SPS-MS3. **E and F**. Experiments performed as described in panels B and C, including a separate analysis of isotopically labeled peptides as single TMT plexes (data labeled as “2XTMT plexes”). For the fractionation experiment, hyperplexed samples were fractionated on stagetip in high pH into two fractions. Average from two individual experiments is plotted.

**Fig 3.**
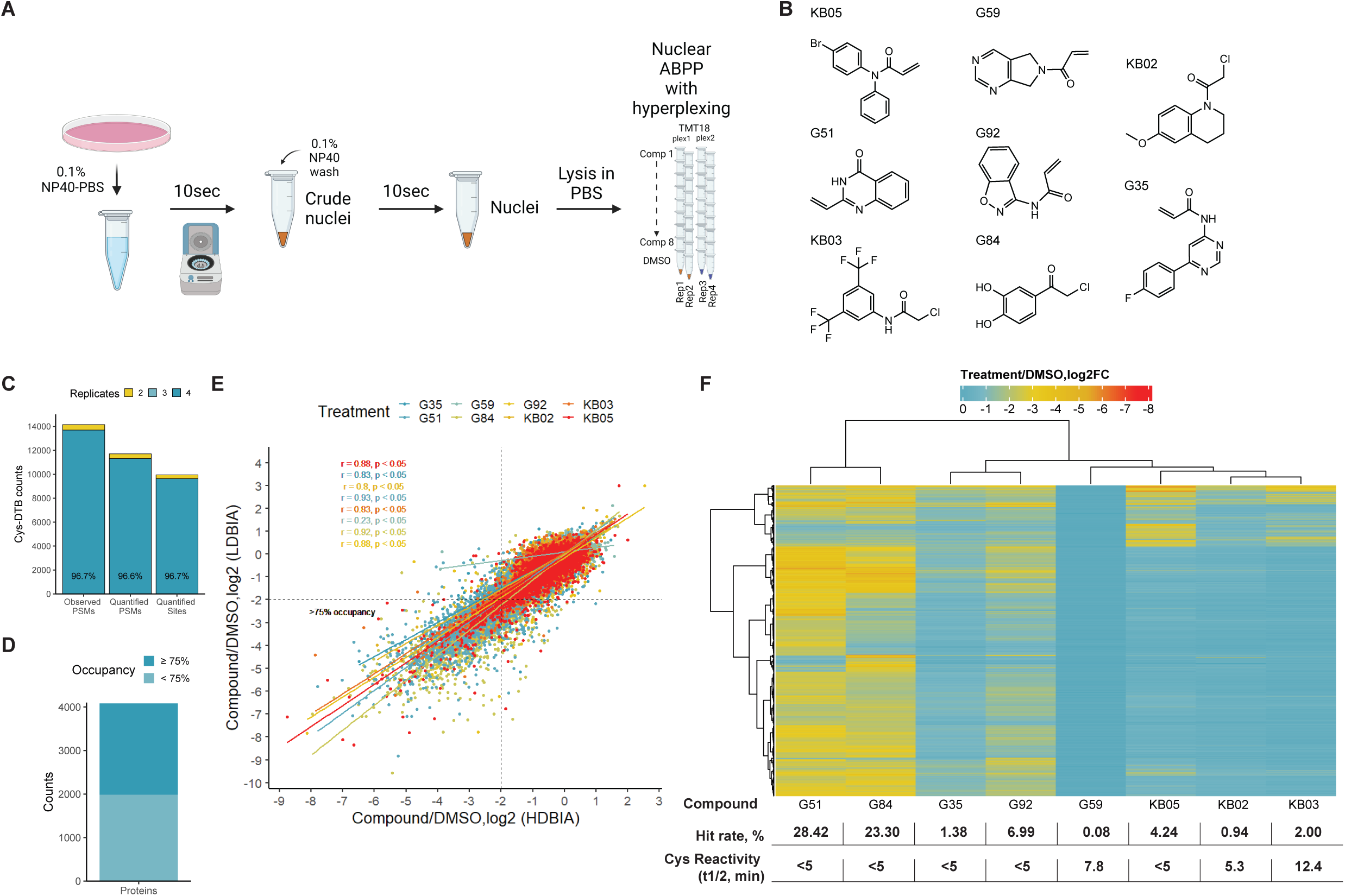
Activity-based proteome profiling in the nuclear fraction. **A**. Cysteine ligandability profiling workflow with HLDBIA/TMT in MCF7 nuclear lysates with electrophilic scout probes (8 treatments, 4 replicates). Nuclear lysates were treated with each compound at 100 µM for 1 hr, followed by HDBIA or LDBIA labeling at 500 µM for 1 hr. **B**. Structures of electrophilic fragments used in the nuclear ABPP experiment. **C**. Number of cysteine sites quantified across two or four replicates in the hyperplexed nuclear ABPP experiment. **D**. Fraction of proteins occupied by at least one electrophilic compound at ≥75% on one or more cysteine sites. **E**. Correlation of replicate measurements derived from “heavy” and “light” biotinylated site abundances as log2 ratios of compound treatment over DMSO control. **F**. Heatmap of site occupancy measured as log2(Treatment/DMSO) across treatments. Summary table includes compound hit rate (fraction of sites occupied by the compound at ≥75%, pvalue≤0.05) and intrinsic compound reactivity measured as the amount of time it takes for 50% of 1 µM compound to be consumed during incubation with 5 mM cysteine (t_1/2_).

First, we tested quantification of paired peptides in a hyperplexed samples using non-RTS and Thermo RTS acquisition methods with Targeted Mass Difference (TMD) trigger. The TMD acquisition parameter triggers MS2 analysis of both isotopically labeled peptide pairs in a hyperplexed sample. As expected, RTS quantification produced an 18% boost in the number of quantified Cys-DTB peptides in SILAC/TMT and HLDBIA/TMT hyperplex analyses over non-RTS method (**Fig. 2B-C**). Interestingly, only a modest 10% improvement or no improvement was observed in the recovery of paired Cys-DTB peptides in SILAC/TMT or HLDBIA/TMT hyperplex, respectively (**Fig. 2B and 2C, top panels**).

Upon closer examination, we observed that some peptides had an isotopic pair that passed online FDR filtering, but not offline FDR filtering. We reasoned that if one PSM from the isotopic pair passed offline FDR filtering, the paired PSM is also a true match to that peptide sequence and should be included in the final peptide identification list. As a result, we observed an additional gain in the number of peptide pairs in the hyperplexed samples (**Figs. 2B and 2C, bottom panels**). However, ∼20-30% of peptides identified in the hyperplexed sample were still missing quantification from their paired isotopic partner.

To address this issue, we developed the PairQuant method within InSeqAPI (**Fig. 2D**). Briefly, using the vendor instrument method, the TMD filter was used to trigger MS/MS analysis of both peptides within a hyperplexed pair. Within inSeqAPI, PairQuant waited for both peptide spectra to be identified before sending instructions to perform separate SPS-MS3 scans on the hyperplexed peptides (**Fig. 2D, 1**). For cases where only one of the hyperplexed peptides was identified due to the correct peptide selection but failure to identify both peptides (**Figure 2D, 2**), TMD selection of incorrect precursor pairs (**Fig 2D, 3**), or MS/MS only performed on one of the hyperplexed peptides within the pair (**Figure 2D, 4**) PairQuant instructed the MS to perform an ion trap MS scan to identify the missing hyperplexed partner. If this scan resulted in the identification of the missing partner peptide then PairQuant instructed the MS to perform separate SPS-MS3 scans on the hyperplexed partners.

As a result, we observed that 98%-100% of peptide pairs had SPS-MS3 scans acquired on both partners when using the PairQuant method (**Figs. 2E and 2F**). While the number of unique cys-DBIA peptides with complete quantitation (i.e., quantified in both hyperplexed pairs) was higher than utilizing two separate TMT plexes, the overall number of unique cys-DBIA peptides was reduced due to the increased sample complexity. To improve depth of cys-DBIA quantification we reduced the complexity of the hyperplexed samples by creating two high-pH fractions through stagetip fractionation (**Figs. 2E and 2F**). These two fractions were analyzed by PairQuant using the same amount of instrument acquisition time as two separate TMT plexes. This resulted in a similar number of quantified unique cys-DBIA peptides when compared to two TMT plexes, but nearly all (98% - 100%) of these were quantified in both of the hyperplexed partner peptides (**Figs. 2E and 2F**). A similar analysis of fractionated hyperplexed samples with Thermo RTS demonstrated a decrease in the total number of quantified unique cys-DBIA peptides as well as the percentage of these peptides that were completely quantified (72% - 78%) (**Figs. 2E and 2F**). These data demonstrate that both the SILAC/TMT and HLDBIA/TMT hyperplexing workflows combined with PairQuant acquisition within InSeqAPI allow for similar depth of peptide identification as a regular TMT workflow, but with a substantial improvement in complete quantification of unique cys-DBIA peptides.

### Hyperplexed cysteine profiling in the nuclear fraction

ABPP with cysteine-targeting covalent fragment compounds has been shown to aid in identification of ligandable pockets across the cellular proteome.^5,10,11^ We wanted to test whether we can improve the throughput and coverage of ligandable pocket profiling in the nucleus by applying an ABPP workflow that included nuclear isolation, hyperplexing, and PairQuant acquisition.

Nuclei were isolated from MCF7 cells using the REAP protocol.^17^ By Western blotting, we observed an enrichment of the nuclear marker histone-H3 and depletion of the cytosolic marker ɑ-tubulin in the nuclear fraction compared to the whole-cell lysate and the cytosolic fraction (**Supp. Fig. 4**). Nuclear lysates in PBS were treated with DMSO or eight cysteine-reactive covalent fragments in four replicates, 36 samples total (**Figs. 3A and 3B**). Upon labeling of unreacted cysteines with LDBIA or HDBIA, “light” and “heavy” sample pairs were combined and processed as a single hyperplex for TMTPro-based quantification using 18 channels. As a result, we were able to quantify 9953 sites from 4081 proteins, with the majority of sites quantified across 4 replicate experiments in the final dataset (**Figs. 3C and 3D, Supp. Table 1**).

For each treatment, measured compound occupancy strongly correlated between HDBIA and LDBIA replicate experiments, except for compound G59, which also showed no significant cys site engagement (**Figs. 3E and 3F, Supp. Table 1**). Although all compounds showed high reactivity in cysteine trapping assay (**Fig. 3F and Supp. Fig. 5**), their promiscuity greatly varied as shown by the heatmap of sites with ≥75% compound occupancy (**Fig. 3F**). Among tested compounds, three, KB02, KB03 and KB05, were previously utilized in other activity-based proteome profiling studies.^5,11,23^ These compounds were shown to have a high coverage of ligandable sites when treatments are done at >100µM concentrations in cell lysates.^5^ Similarly, we observed ∼1-4% site labeling by these compounds in the nuclear fraction. Two compounds with even greater promiscuity of 23% and 28% were a chloroacetamide compound G84 and a quinazolinone-based compound G51, respectively (**Fig. 4F**). These results demonstrate that the hyperplexing approach allows for fast and accurate screening of compounds with various reactivity across the nuclear proteome.

**Fig 4.**
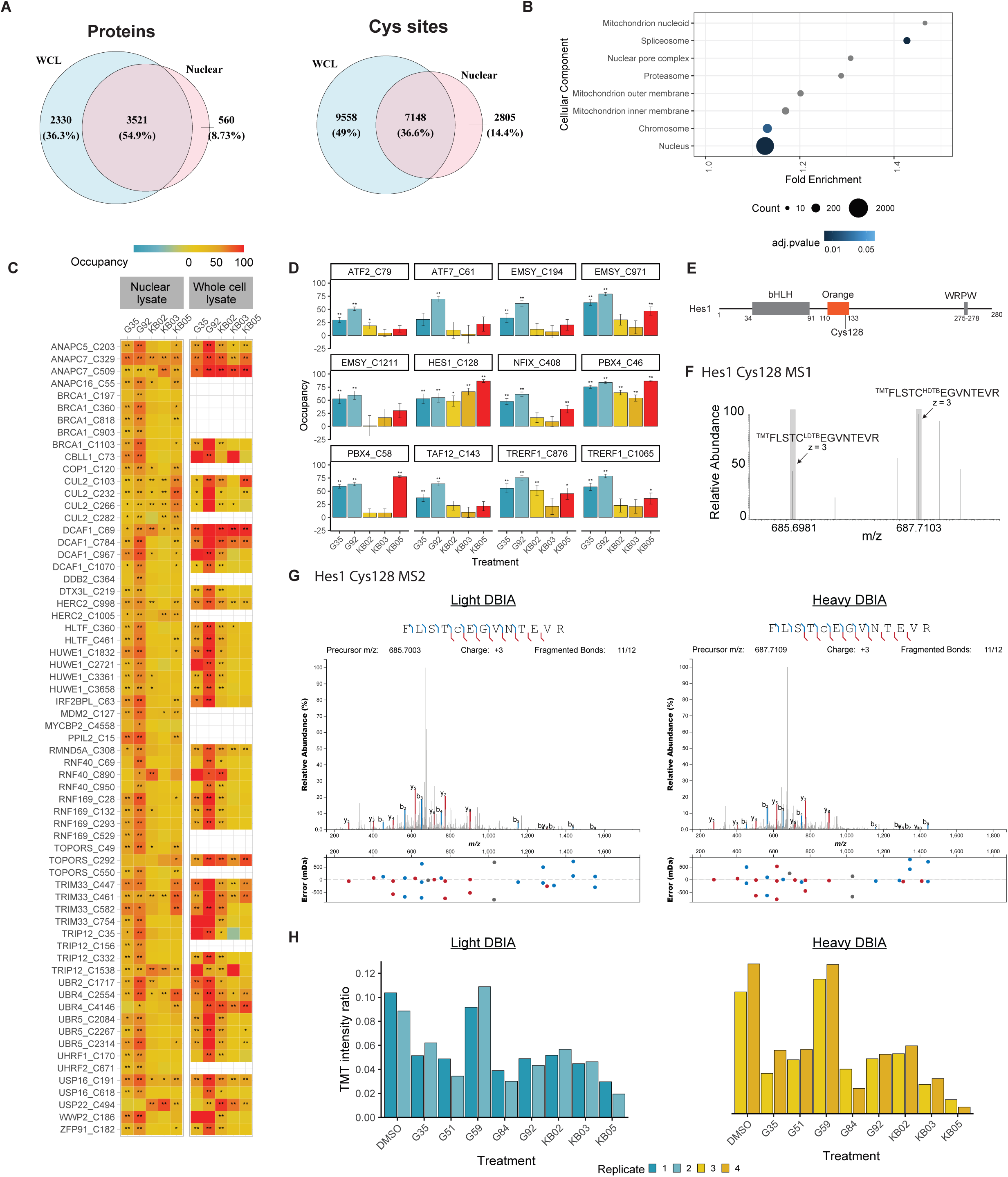
Characterization of ligandable cysteine sites in the nuclear proteome. **A**. For the whole-cell lysate ABPP experiment, MCF7 lysates were incubated with 500µM of each electrophilic compound for 1 hr, followed by labeling with 500 µM LDBIA probe for 1 hr. Nuclear lysates were treated with each compound at 100 µM for 1 hr as described in Fig. 3. Venn diagrams showing the number of proteins (left panel) and unique cysteine sites (right panel) identified in each experiment. **B**. Gene Ontology Cellular Component analysis of proteins identified in the nuclear fractions. **C**. Heatmaps of site occupancies of proteins in ubiquitination pathway observed in the nuclear and whole-cell ABPP experiment with indicated electrophilic fragments. **D**. Site occupancy of selected transcription regulators uniquely identified in the nuclear ABPP experiment. **E**. Localization of Cys128 in the “Orange” domain of HES1 transcription repressor. **F** and **G**. MS1 and MS2 spectra of HES1 Cys128 peptide modified with HDBIA or LDBIA. **H**. Relative abundance of HES1 Cys128 peptide modified with HDBIA or LDBIA between conditions.

### Ligandable nuclear targets

To compare the depth of nuclear cysteine peptide enrichment to a whole-proteome profiling experiment, we performed a standard ABPP experiment in whole cell lysates from MCF7 cells. InSeqAPI RTS acquisition of biotinylated cysteine peptides from whole-cell lysates resulted in quantification of 5851 proteins with 16706 cysteine sites from three TMT plexes (**Supp. Table 2**). We observed that 2805 sites from 1784 proteins were uniquely identified in the nuclear fraction compared to the whole-cell lysate; this included 560 proteins that were quantified only in the nuclear experiment (**Fig. 4A**). The cellular component annotation analysis of the proteins detected in the nuclear fraction revealed significant enrichment of the nucleus-localized proteins (**Fig. 4B, Supp. Table 1**), supporting our hypothesis that we can improve the depth of activity-based proteome profiling studies through subcellular fractionation.

For further evaluation of ligandable nuclear cysteine sites, we focused on the compounds that showed medium promiscuity in ABPP experiment, KB02, KB03, KB05, G35 and G92 (**Fig. 3F**). Among liganded proteins in the nuclear fraction (i.e. occupancy ≥50%), we observed 21 E3 ubiquitin-protein ligases and core components of their complexes (such as Skp, Cullin, F-box containing complex, SCF), as well as two deubiquitinases (DUBs) (**Fig. 4C, Supp. Table 1**). These proteins localize and function primarily or partially in the nucleus, participating in processes such as DNA damage response (BRCA1, HERC2, RNF169, TRIP12, HUWE1, UBR5) and transcription (TRIM33, IRF2BPL, UBR2, USP16, USP22, DCAF1, HUWE1, RNF40). DUB enzymes ubiquitin carboxyl-terminal hydrolase 16 and 22 (USP16 and USP22) function in the transcription regulation in the nucleus through deubiquitination of histone substrates (reviewed in^24^). USP22 Cys 494 (87.5% occupancy by KB03) is localized in the USP domain,^25^ and USP16/Ubp-M Cys 191 (78% occupancy by G92) is adjacent to the USP domain.^26^ Both of these sites were also liganded in the whole-cell lysate analysis, further supporting the finding of the nuclear fractionation experiment (**Fig. 4C**). We also observed ligandable sites on three E3 ubiquitin ligases, UBR2, UBR4 and UBR5 that function in the N-end rule proteolytic pathway. Their targets are usually short-lived proteins that are recognized by n-degrons that consist of exposed or modified n-terminal residues.^27^ UBR2 has an essential function in regulation of chromatin-wide transcriptional silencing through ubiquitination of H2A histone.^28^ UBR2 Cys1717 was liganded by G92 in the nuclear and whole-cell lysates, and is localized in the UBR autoinhibitory domain (UAIN) at the protein c-terminus.^29^ A few sites were uniquely quantified in the nuclear fraction, such as BRCA1 C360. This site is present in the BRCA1 region interacting with DNA recombinase RAD50.^30^ Cys360 was liganded by G92 and G35 electrophiles at 73% and 53% occupancy, respectively. Therefore, nuclear activity-based proteome profiling allows for identification of new sites in the protein-protein interaction domains of nuclear ubiquitin ligases and deubiquitinases.

Among 577 proteins unique to the nuclear fraction, we identified 27 transcription regulators (**Supp. Table 1**). Transcription factors represent challenging targets for drug development, but their function can potentially be modulated through protein-protein and protein-DNA interactions. In the nuclear ABPP analysis, we identified ligandable cysteine sites on 12 transcription regulators (**Fig. 4D**). ATF2 Cys79 is localized in a zinc-coordinated transactivation domain and was liganded by G92 at 50.8%. Interestingly, another activation transcription factor family member (ATF7) was also liganded by G92 at a homologous site. These cysteine sites are localized near a phosphoswitch motif regulated by MAPK-mediated phosphorylation.^31^ Transcription activator PBX4 was occupied at 59-77% by three fragments, G35, G92 and KB05 on Cys 58, which is present in protein-protein interaction domain PBC-A.^32^ Transcriptional regulating protein of 132 kDa (TReP-132/TRERF1) was liganded by four compounds at 50-79% on Cys 876, which is localized in SANT domain that may mediate protein-protein interactions with HDAC1/2 within MiDAC complex.^33^ Transcription factor HES1 was liganded by all compounds with medium promiscuity at 48-87% occupancy on Cys 128. HES1 is a basic helix-loop-helix (bHLH) transcription repressor that contains a unique Orange domain with unknown function, which is present exclusively in transcription repressors (**Fig. 4E**).^34^ Quantification of Cys 128 abundance was based on detection of the light and heavy Cys-DTB pair of ions (**Fig. 4F**), both of which were selected for MS2 sequencing. Eight TMT-containing b ion fragments from each MS3 spectrum were assigned by InSeqAPI for MS3 fragmentation (**Fig. 4G**). Relative TMT reporter ion signal was reproducible between heavy and light-labeled peptide species and within each TMT plex (**Fig. 4H**). Therefore, our hyperplexing approach enabled by PairQuant acquisition method can provide accurate quantification of cysteine engagement in activity-based proteome profiling studies by reducing the number of missing values between replicate TMT plexes.

## Discussion

In this work, we described the analysis of ABPP samples using intelligent data acquisition (IDA) enabled by InSeqAPI. We focused specifically on improving sample throughput by employing hyperplexed analysis, an approach where samples are isotopically labeled on protein level for replicate analysis before TMT labeling at the peptide level to enable comparisons between conditions. To achieve complete coverage of hyperplexed peptide pairs we introduced PairQuant, a method that utilizes a real-time database search to trigger quantitative scans only when both hyperplexed peptide pairs have been identified. If only one of the peptides within a hyperplexed pair is identified, PairQuant sends an identification scan for the missing hyperplexed partner. This approach improved the quantification of hyperplexed pairs to ∼98- 100% after rescuing partners that were matched to the correct peptide, but did not pass offline FDR filters. A challenge of this approach is that increased sample complexity led to a decrease in the total number of unique cysteine containing peptides quantified in a single run. To reduce sample complexity, hyperplexed samples were fractionated into two fractions and analyzed in the same amount of instrument time as two-plex TMT analysis. We demonstrated that this approach allows for a significant reduction in missing values between plexes (<2%), while maintaining the same number of total unique peptide sequences quantified.

While our analyses featured two hyperplexed TMT plexes, the level of hyperplexing can be increased and potentially incorporated into specific workflows through the use of alternative metabolic labeling strategies. For instance, triple SILAC would allow for triplicate analysis of 18 conditions (54 samples total) in an ABPP workflow. Such labeling approach was previously demonstrated in combination with a 6-plex TMT to monitor yeast proteome changes in response to rapamycin stimulation. ^15^ Additionally, the use of metabolic labeling by isotopic tyrosine was previously demonstrated in phosphoprofiling studies.^35^ Therefore, incorporation of tyrosine SILAC into a hyperplexed method could extend the number of timepoints used in phosphosignaling studies.

We also tested ABPP with a nuclear enrichment strategy, where nuclear extracts were labeled with scout probes, promiscuous electrophilic fragments with highly reactive cysteine-targeting warheads. A single hyperplexed nuclear sample combined 36 conditions which included eight compounds and a DMSO control in four replicates, where two replicates were labeled with isotopically “light” and “heavy” cysteine-reactive tags and two replicates were isobarically tagged by an 18-plex TMT reagent. Although all of the scout fragments tested were intrinsically highly reactive and stable in PBS over the labeling period, we observed a broad range of hit rates across compounds. This result underlies the complexity of electrophilic compound interactions with their targets that is not driven purely by the warhead reactivity. One of the advantages of nuclear enrichment strategy in the covalent fragment screening is the removal of highly reactive cysteine sites present in the cytosolic and mitochondrial proteome, which can outcompete less reactive sites in the nuclear proteome. As a result, nuclear enrichment allowed for identification of ligandable cysteine sites on nuclear-localized proteins that were missed in a whole-proteome analysis, including nuclear ubiquitin ligase machinery and transcription regulators.

Recent studies have shown that covalent ligands can be used as molecular glues^36^ or can be incorporated into heterobifunctional electrophilic proteolysis targeting chimeras (PROTACs) for targeted degradation.^23,37–39^ As such, covalent small molecules also provide opportunities for targeting functions of transcription factors, which are historically challenging to drug. For instance, identification of a ligandable cysteine within the Orange domain of HES1 transcription repressor highlights a possibility for developing a covalent small molecule that modulates its functions through protein-protein interactions and/or a covalent PROTAC.

In summary, this study describes an application of intelligent data acquisition in hyperplexed activity-based proteome profiling studies, which increases experimental throughput and completeness of the site occupancy data. Importantly, the PairQuant data acquisition method can be used on any hyperplexed sample, potentially unlocking this approach for other proteomics applications. For example, hyperplexing was recently described for single cell proteomics in a technique termed “hyperSCP”, here multiplexing was increased to 28 samples per run making it possible to analyze up to ∼280 cells per day.^40^ Lastly, this work represents an introduction inSeqAPI, an iAPI framework that enables rapid development of custom IDA methods. While this study focused on the PairQuant method, inSeqAPI will continue to evolve with methods tailored specifically for other proteomics approaches (e.g., single cell proteomics, HLA peptidomics, etc.).

## Supporting information

SuppTable1

SuppTable2

## Acknowledgements

The authors thank current and former members of the MPL+NGS department for their input and support, former and current members of Discovery Chemistry department for their help with covalent probe synthesis.

## Author contributions

1. H. G. B., and C. M. R. conceptualization; H. G. B. and D. S. K. methodology; H. G. B., T. P. M., S. W., M. C., C. M. R. formal analysis; H. G. B., T. P. M., S. W., M. C., C. M. R. investigation; H. G. B., and C. M. R. writing–original draft; H. G. B., T. P. M., S. W., M. C., C. M. R. writing– review and editing; C. M. R. supervision.

---

**Supplemental Fig.1.**
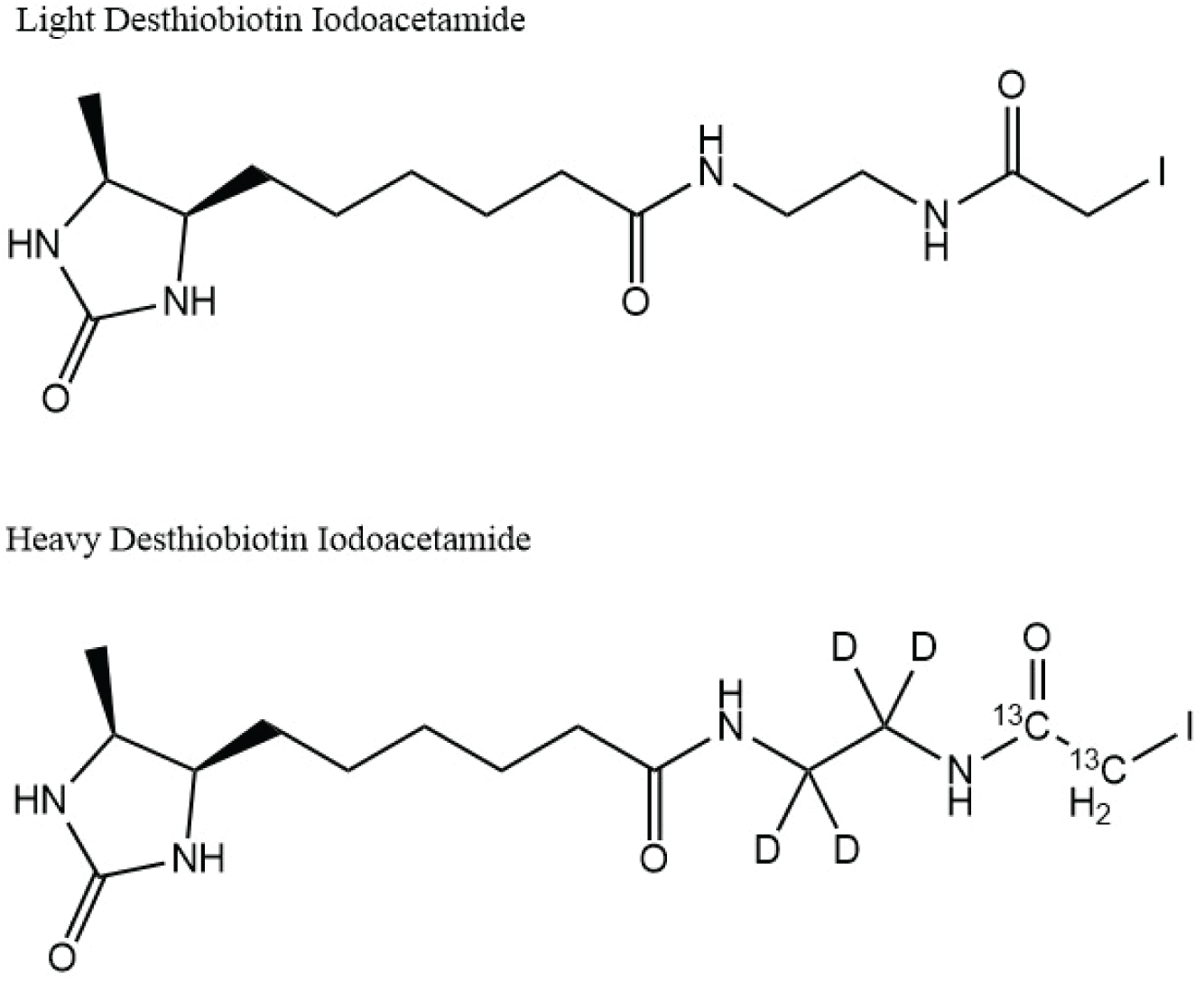
Structures of LDBIA and HDBIA reactive probes.

**Supplemental Fig.2.**
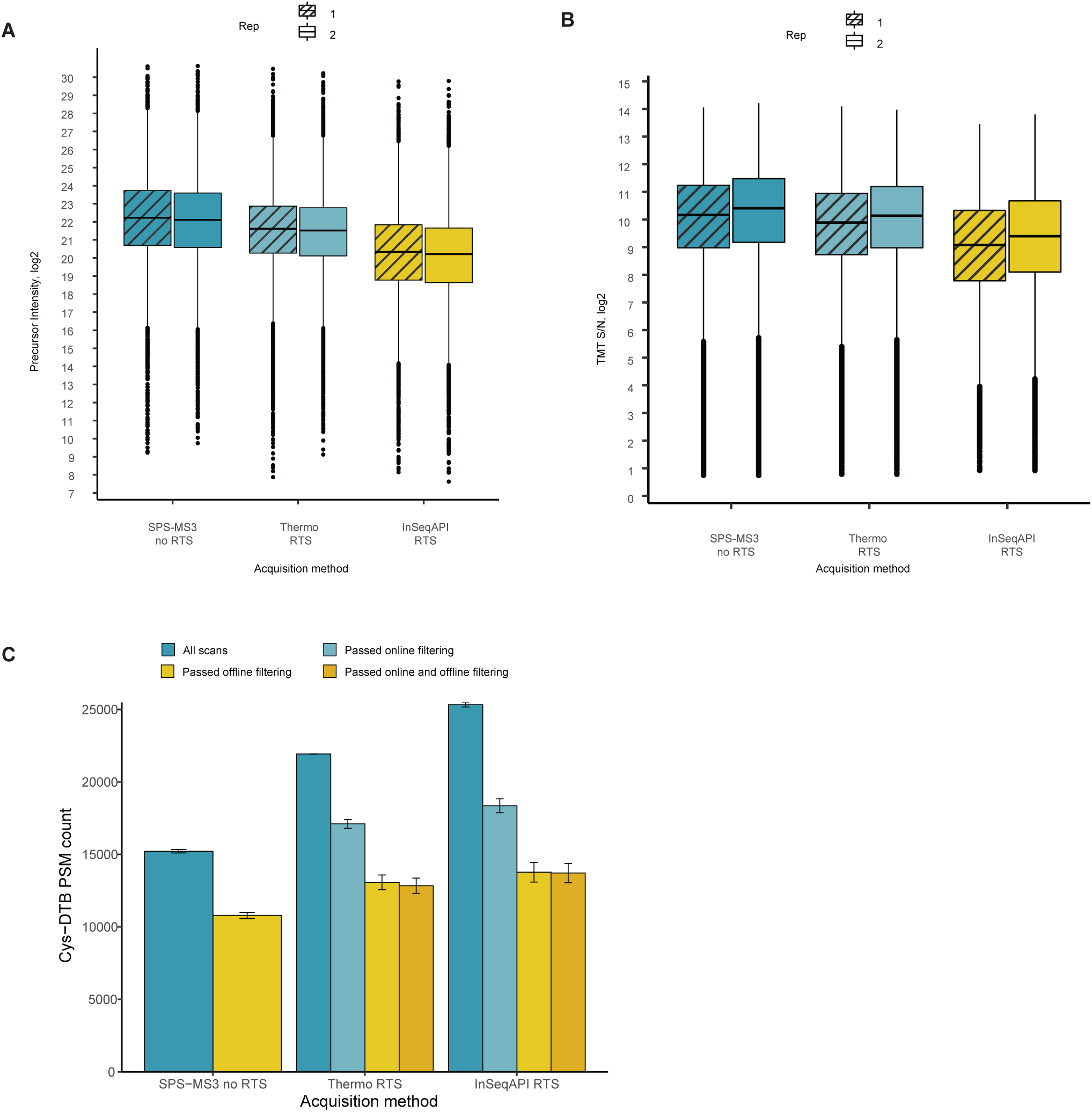
**A**. and **B**. Precursor ion intensity of biotinylated cys peptides (A) and TMT S/N distribution from all MS3 spectra (B) between runs acquired with SPS-MS3 without RTS and with indicated RTS method (two individual experiments), data from experiment in Fig. 1C. **C**. Number of biotinylated cysteine PSMs identified using indicated acquisition method before peptide FDR filter (“All scans”), after RTS peptide FDR filter (“Passed online filtering”), after offline search and peptide FDR filter (“Passed offline filtering”), and after RTS and offline FDR filter (“Passed online and offline filtering”). Data from experiment in Fig. 1C.

**Supplemental Fig.3.**
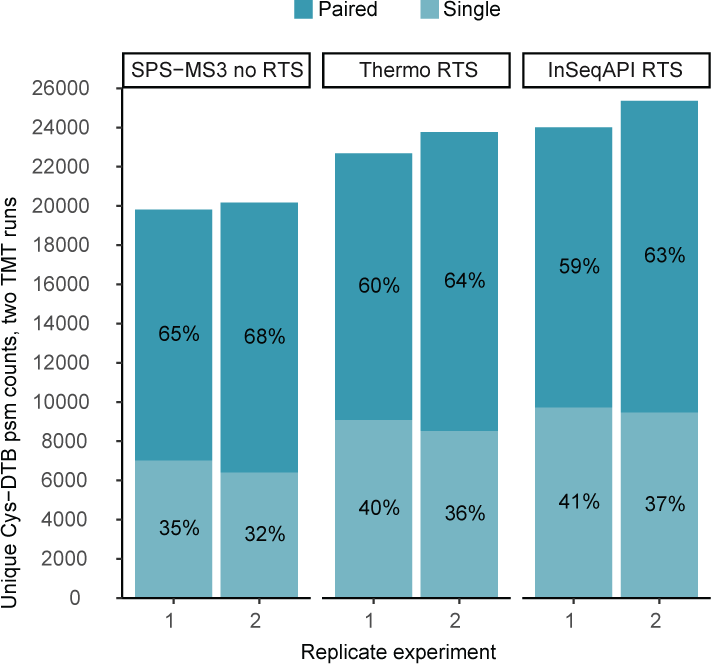
Lysates from C2C12 “heavy” and “light” SILAC cell culture were labeled with 500 µM LDBIA probe for 1 hr. After digestion and streptavidin enrichment, biotinylated peptides were analyzed using indicated acquisition methods. Number of biotinylated cysteine PSMs identified between two TMT runs (“heavy” SILAC and “light” SILAC) is shown from two individual experiments.

**Supplemental Fig.4.**
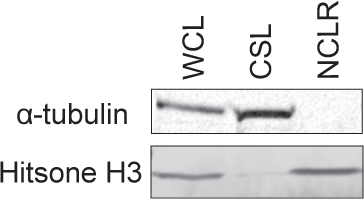
MCF7 cells were fractionated into nuclear and cytosolic fraction using REAP protocol^17^. Fractions were analyzed by Western Blotting with anti-tubulin and anti-histone H3 antibodies as cytosolic and nuclear markers, respectively.

**Supplemental Fig.5.**
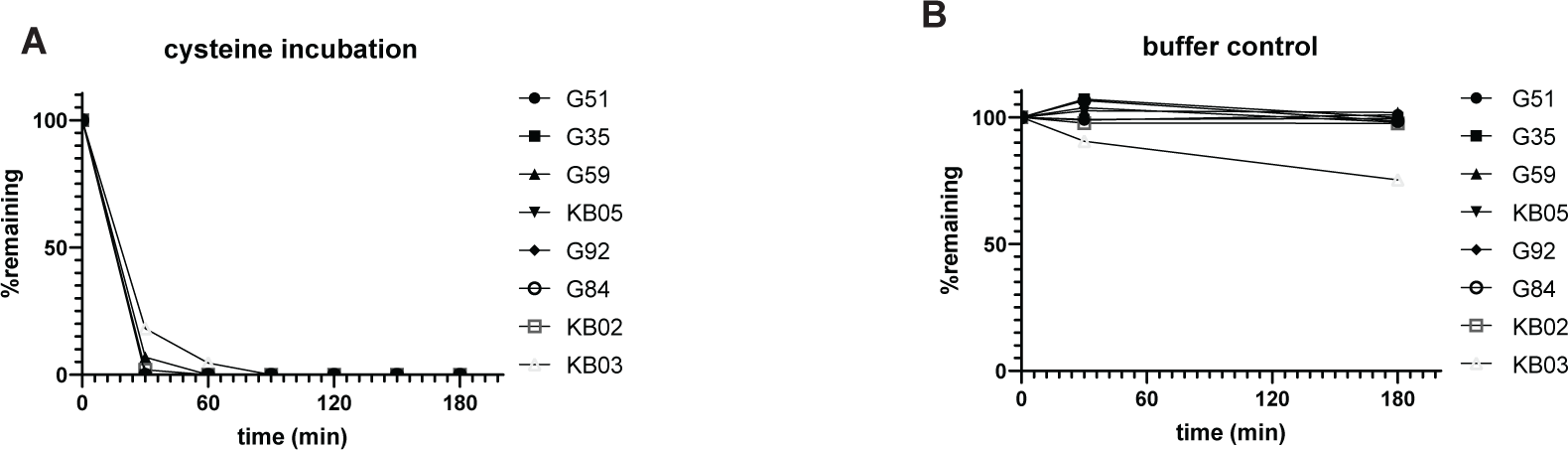
**A**. Test compounds (1 µM) and cysteine (5 mM) were mixed sequentially to start the reaction. Aliquots were taken at indicated timepoints and quantified via LC-MS/MS. Percentage remaining was calculated using peak area ratios normalized to the 0 minutes time-point sample (% *remaining* = 100 ∗ *e*^−*kreact* ∗ 5 *mM* ∗ *time*^). **B**. Incubation without cysteine was also carried out to check compound stability in the buffer.

**Supplemental Fig.6.**
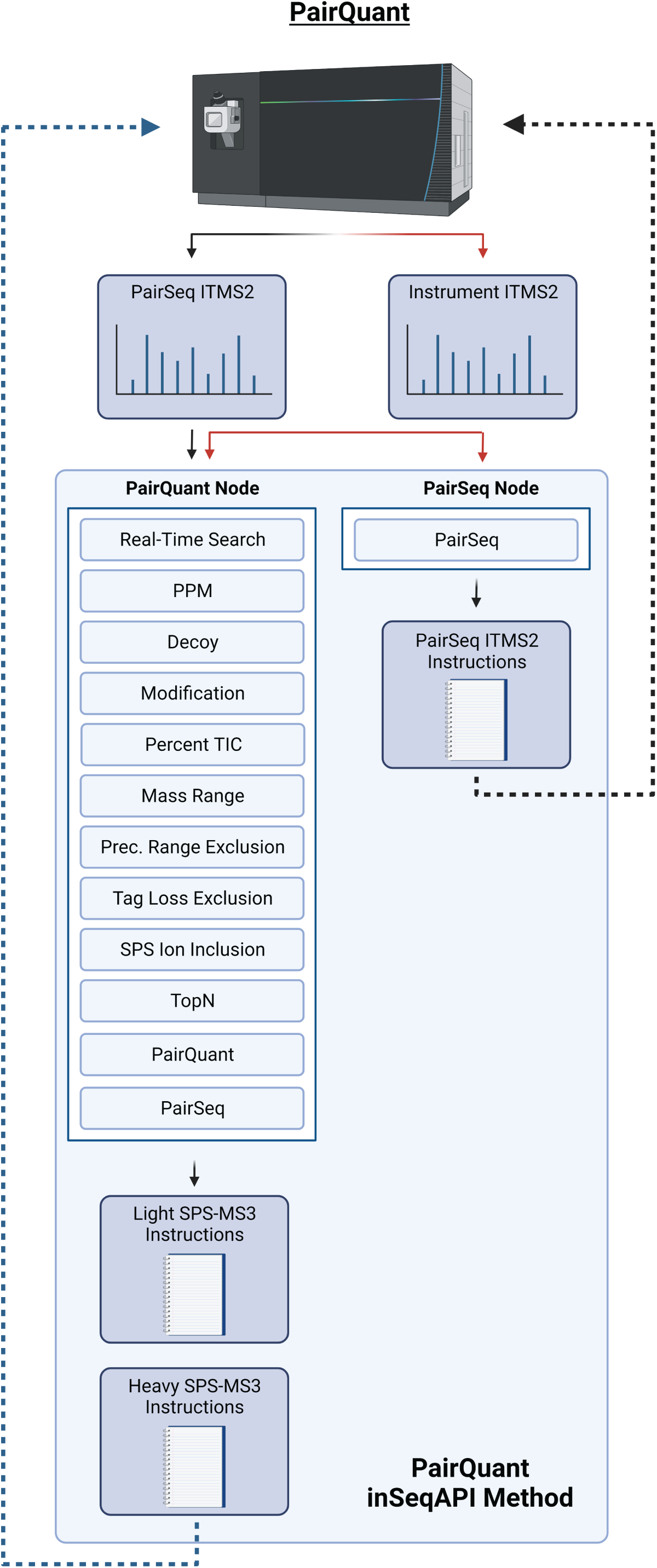
Detailed schematic PairQuant method within inSeqAPI.

